# Through a Dog’s Eyes: fMRI Decoding of Naturalistic Videos from Dog Cortex

**DOI:** 10.1101/2022.07.12.499776

**Authors:** Erin M. Phillips, Kirsten D. Gillette, Daniel D. Dilks, Gregory S. Berns

## Abstract

Recent advancements using machine learning and fMRI to decode visual stimuli from human and nonhuman cortex have resulted in new insights into the nature of perception. However, this approach has yet to be applied substantially to animals other than primates, raising questions about the nature of such representations across the animal kingdom. Here, we used awake fMRI in two domestic dogs and two humans, obtained while each watched specially created dog-appropriate naturalistic videos. We then trained a neural net (Ivis) to classify the video content from a total of 90 minutes of recorded brain activity from each. We tested both an object-based classifier, attempting to discriminate categories such as dog, human and car, and an action-based classifier, attempting to discriminate categories such as eating, sniffing and talking. Compared to the two human subjects, for whom both types of classifier performed well above chance, only action-based classifiers were successful in decoding video content from the dogs. These results demonstrate the first known application of machine learning to decode naturalistic videos from the brain of a carnivore and suggest that the dog’s-eye view of the world may be quite different than our own.

## Introduction

The brains of humans, like other primates, demonstrate the parcellation of the visual stream into dorsal and ventral pathways with distinct, and well-known, functions – the ‘what’ and ‘where’ of objects ^1^. This what/where dichotomy has been a useful heuristic for decades, but its anatomical basis is now known to be much more complex, with many researchers favoring a parcellation based on recognition versus action (‘what’ versus ‘how’) ^2–5^. And while our understanding of the organization of the primate visual system continues to be refined and debated, much remains unknown about how the brains of other mammalian species represent visual information. In part, this lacuna is a result of the historical focus on a handful of species in visual neuroscience. New approaches to brain imaging, however, are opening up the possibility of noninvasively studying the visual systems of a wider range of animals, which may yield new insights into the organization of the mammalian nervous system.

Dogs (*Canis lupus familiaris*) present a rich opportunity to study the representation of visual stimuli in a species evolutionarily distant from primates, as they may be the only animal that can be trained to cooperatively participate in MRI scanning without the need for sedation or restraints ^6–8^. Because of their co-evolution with humans over the last 15,000 years, dogs also inhabit our environments and are exposed to many of the stimuli that humans encounter on a daily basis, including video screens, which are the preferred way of presenting stimuli in an MRI scanner. Even so, dogs may process these common environmental stimuli in ways that are quite different than humans, which begs the question of how their visual cortex is organized. Basic differences – such as a lack of a fovea, or being a dichromat – may have significant downstream consequences not only for lower-level visual perception but also for higher-level visual representation. Several fMRI studies in dogs have demonstrated the existence of both face- and object-processing regions that appear to follow the general dorsal/ventral stream architecture seen in primates, although it remains unclear whether dogs have face-processing regions per se, or whether these regions are selective for the morphology of the head, e.g. dog versus human ^9–13^. Regardless, the brain of a dog, being smaller than most primates, would be predicted to be less modularized ^14^, so there may be more intermixing of types of information in the streams, or even privileging of certain types of information, like actions. It has been suggested, for example, that movement might be a more salient feature in canine visual perception than texture or color ^15^. Additionally, as dogs do not have hands, one of the primary means through which we interact with the world, their visual processing, particularly of objects, may be quite different than primates’. In line with this, we recently found evidence that interaction with objects by mouth, versus paw, resulted in greater activation in object-selective regions in the dog brain ^16^.

Although dogs may be accustomed to video screens in their home environment, that does not mean they are used to looking at images in an experimental setting the same way a human would. The use of more naturalistic stimuli may help to resolve some of these questions. In the last decade, machine learning algorithms have achieved considerable success in decoding naturalistic visual stimuli from human brain activity. Early successes focused on adapting classical, blocked, designs to use brain activity to both classify the types of stimuli an individual was seeing, as well as the brain networks that encoded these representations ^17–19^. As more powerful algorithms were developed, especially neural networks, more complex stimuli could be decoded, including naturalistic videos ^20,21^. These classifiers, which are typically trained on neural responses to these videos, generalize to novel stimuli, allowing them to identify what a particular subject was observing at the time of the fMRI response. For example, certain types of actions in movies can be accurately decoded from the human brain, like jumping and turning, while others, e.g. dragging, cannot ^22^. Similarly, although many types of objects can be decoded from fMRI responses, general categories appear to be more difficult. Brain decoding is not limited to humans, providing a powerful tool to understand how information is organized in the brains of other species. Analogous fMRI experiments with nonhuman primates have found distinct representations in the temporal lobe for dimensions of animacy and faciness/bodiness, which parallels that in humans ^23^.

As a first step toward understanding dogs’ representation of naturalistic visual stimuli, we used awake fMRI in two highly MRI-adept domestic dogs to measure cortical responses to dog-appropriate videos. We used naturalistic videos because of their potentially greater ecological validity to a dog and because of their demonstrated success with neural nets that map video content to dog movement ^24^. Over three separate sessions, we obtained 90 minutes of fMRI data from each dog’s response to 256 unique video clips. For comparison, we did the same for two human volunteers. Then, using a neural network, we trained and tested classifiers to discriminate either ‘objects’ (e.g. human, dog, car) or ‘actions’ (e.g. talking, eating, sniffing) using varying numbers of classes. Our goals were two-fold: 1) determine whether naturalistic video stimuli could be decoded from dog cortex; and 2) if so, provide a first look into whether the organization was similar to that of humans.

## Results

To different degrees, the neural net classifier was successful for both humans and dogs. For humans, the algorithm was able to classify both objects and actions, with three-class models for both achieving a mean accuracy of 70%. We used the Label Ranking Average Precision (LRAP) score as the primary metric to compute the accuracy of the model on the test set. This metric measures the extent to which the classifier assigned higher probabilities to true labels ^25^. For both humans, the median LRAP scores were greater than the 99th percentile of a randomly permuted set of labels for all models tested (Table 1; Fig. 2). For dogs, only the action model had a median LRAP percentile rank significantly greater than chance in both participants (Table 1; p=0.13 for objects and p<0.001 for actions; mean three-class action model LRAP score for dogs = 78^th^ percentile). These results were true for all subjects individually as well as when grouped by species.

**Table 1:** Aggregated metrics of the Ivis machine learning algorithm over 100 iterations of training and testing on BOLD responses to naturalistic video stimuli obtained via awake fMRI in dogs and humans. Object models had three target classes (’dog’, ‘human’, ‘car’) and action models had either three or five classes (three class: ‘talking’, ‘eating’, ‘sniffing’; five class: ‘talking’, ‘eating’, ‘sniffing’, ‘petting’, ‘playing’). Values significantly greater than chance are shown in bold.

|  | Model Type | Training Accuracy | Test Accuracy | F1 Score | Precision | Recall | LRAP score median percentile |
| --- | --- | --- | --- | --- | --- | --- | --- |
| Human 1 | Object (3 class) | 0.98 | 0.69 | 0.48 | 0.52 | 0.49 | **>99** |
|  | Action (3 class) | 0.98 | 0.72 | 0.51 | 0.54 | 0.54 | **>99** |
|  | Action (5 class) | 0.97 | 0.51 | 0.28 | 0.37 | 0.27 | **>99** |
| Human 2 | Object (3 class) | 0.98 | 0.68 | 0.45 | 0.50 | 0.47 | **>99** |
|  | Action (3 class) | 0.98 | 0.69 | 0.46 | 0.50 | 0.48 | **>99** |
|  | Action (5 class) | 0.97 | 0.53 | 0.30 | 0.40 | 0.27 | **>99** |
| Bhubo | Object (3 class) | 0.99 | 0.61 | 0.38 | 0.41 | 0.39 | 57 |
|  | Action (3 class) | 0.98 | 0.63 | 0.38 | 0.40 | 0.40 | **87** |
|  | Action (5 class) | 0.99 | 0.45 | 0.16 | 0.29 | 0.13 | **88** |
| Daisy | Object (3 class) | 1.00 | 0.61 | 0.38 | 0.43 | 0.39 | 43 |
|  | Action (3 class) | 0.97 | 0.62 | 0.35 | 0.38 | 0.35 | **60** |
|  | Action (5 class) | 0.99 | 0.44 | 0.16 | 0.27 | 0.13 | **76** |

**Figure 1.**
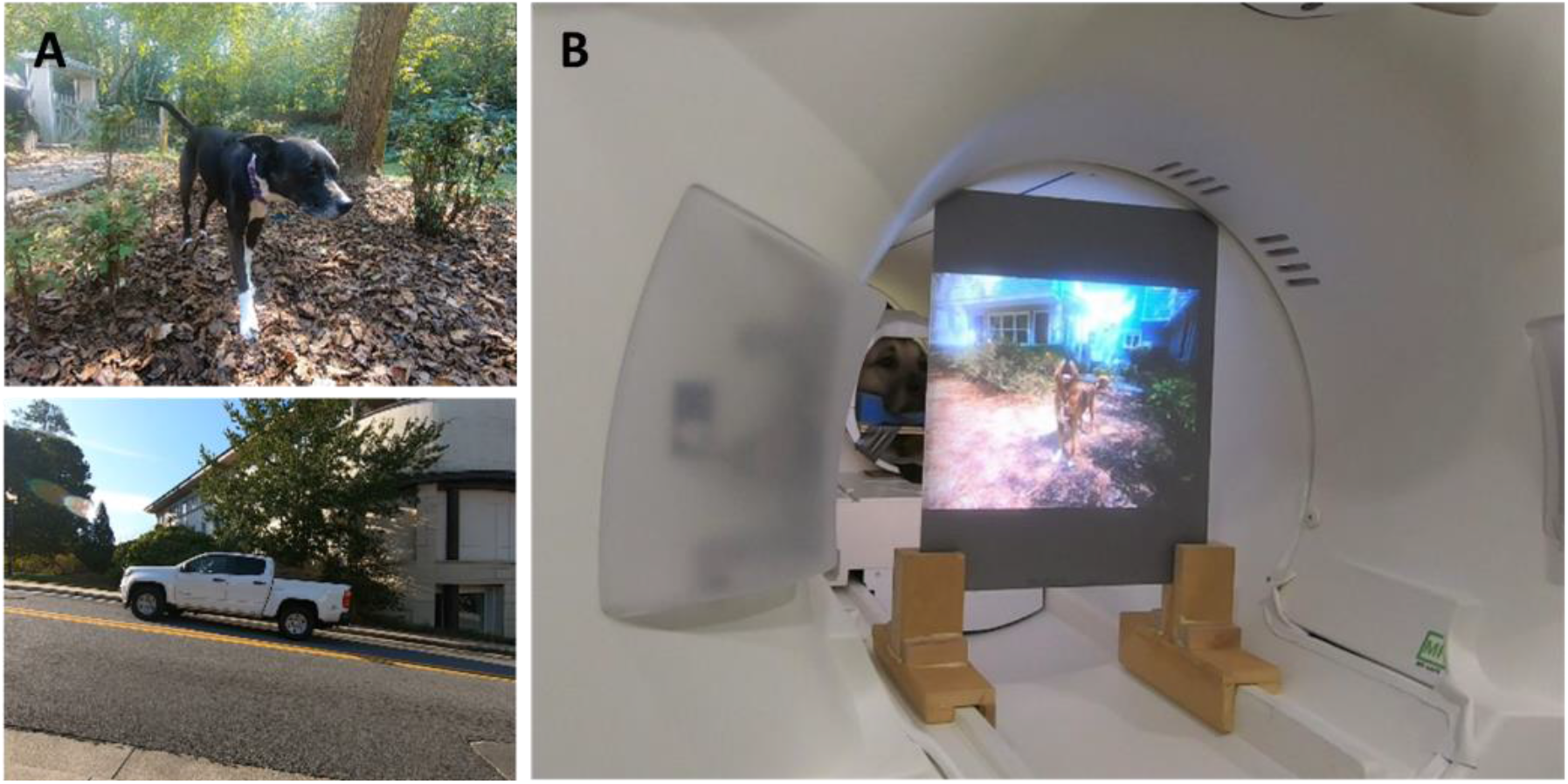
Naturalistic videos and presentation in MRI bore. **A)** Example frames from video clips shown to participants. **B)** Bhubo, a 4-year-old Boxer-mix, watching videos while undergoing awake fMRI.

**Figure 2.**
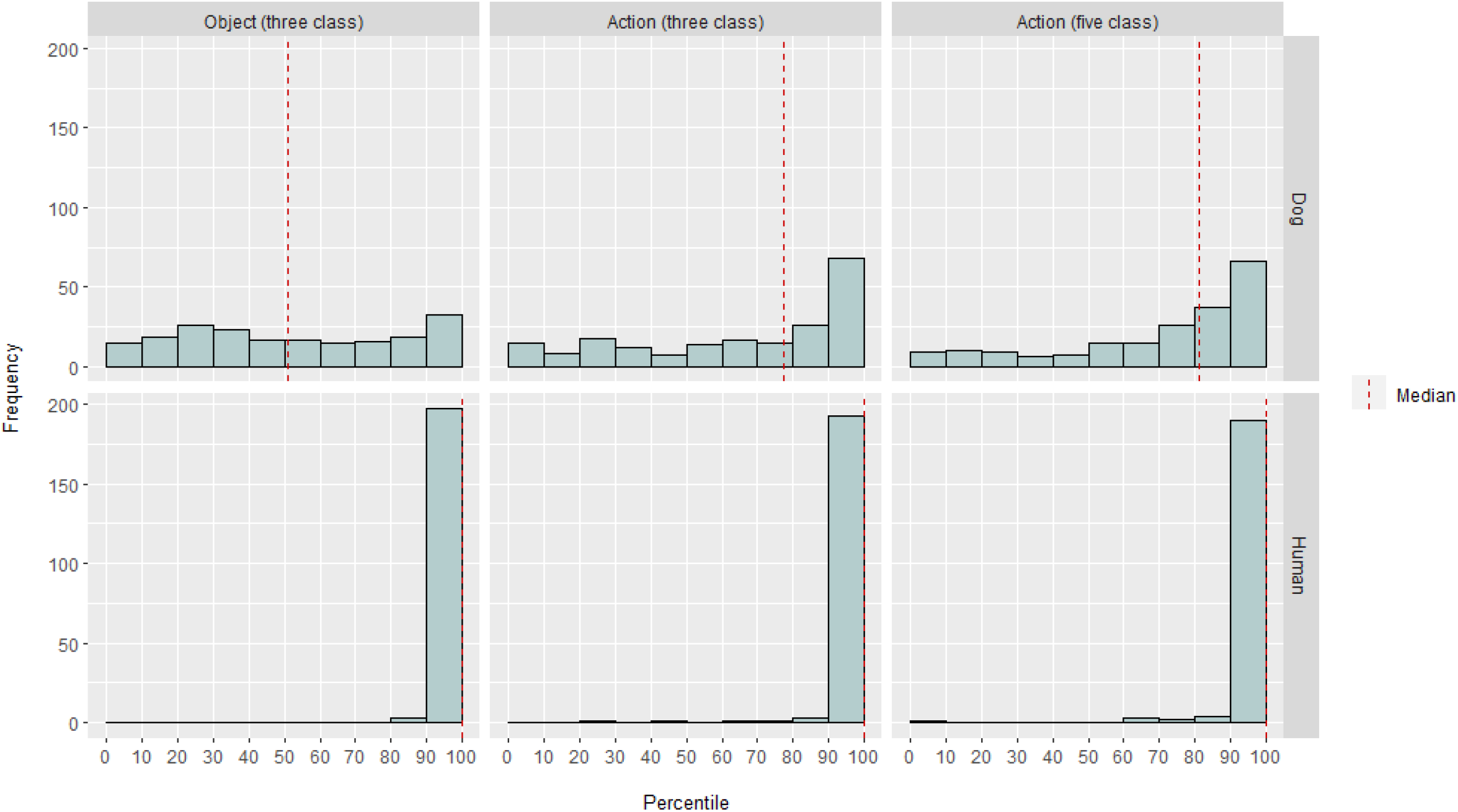
Model performance in dogs and humans. The distribution of LRAP scores, presented as percentile rankings of their null distributions, over 100 iterations of training and testing the Ivis machine learning algorithm for a three-class object-based model, a three-class action-based model and a five-class action-based model, where models attempted to classify BOLD responses to naturalistic video stimuli obtained via awake fMRI in dogs and humans. Scores are aggregated by species. An LRAP score with a very high percentile ranking indicates that the model would be very unlikely to achieve that LRAP score by chance. A model performing no better than chance would have a median LRAP score percentile ranking of ∼50. Dashed lines represent the median LRAP score percentile ranking for each species across all 100 runs.

Given the classifier’s success, we trained and tested with additional classes to determine the limits of the model. Of the additional models tested, the model with the highest median LRAP percentile ranking in both dogs had five classes: the original ‘talking’, ‘eating’ and ‘sniffing’, plus two new classes, ‘petting’ and ‘playing’ (Fig. 2). This model had a median LRAP percentile rank significantly greater than that predicted by chance for all participants (Table 1; p<0.001 for both dogs and humans; mean five-class action model LRAP score for dogs = 81^st^ percentile).

When back-mapped to their respective brain atlases, feature importance scores of voxels revealed a number of clusters of informative voxels in the occipital, parietal, and temporal cortices of both dogs and humans (Fig. 3). In the humans, the object-based and action-based models revealed a more focal pattern than in the dogs, and in regions typically associated with object recognition, although with slight differences in spatial location of object-based voxels and action-based voxels.

**Figure 3.**
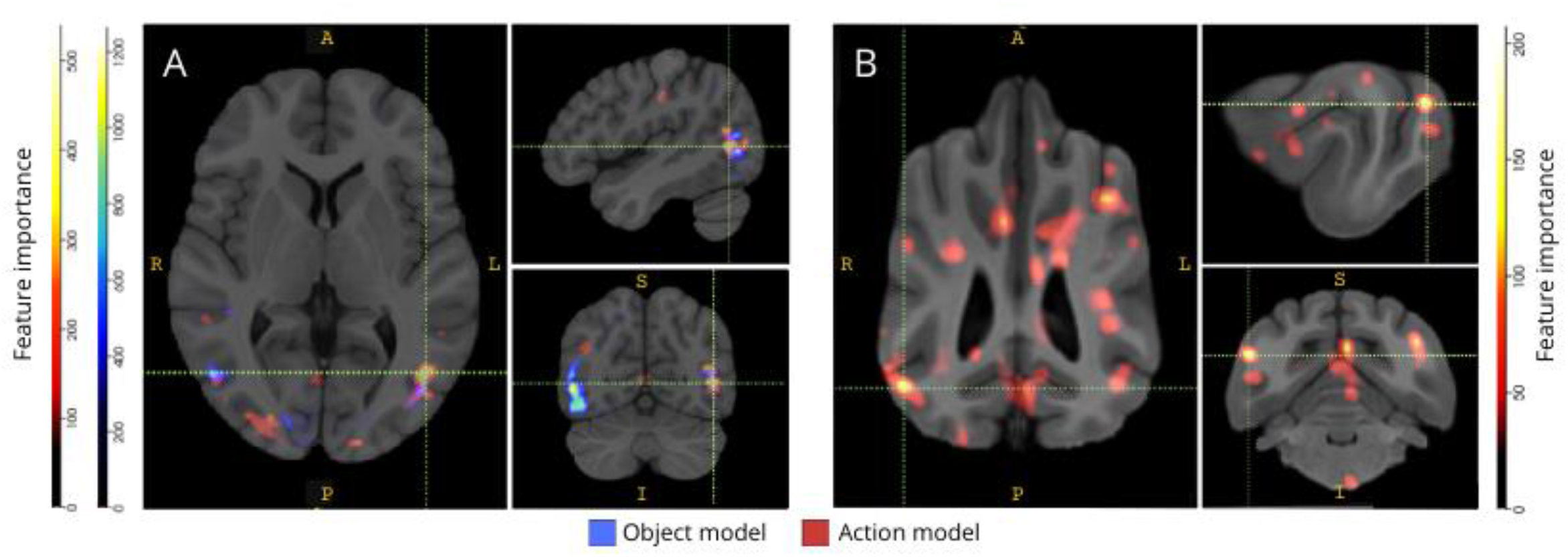
Regions important for the discrimination of three-class object and five-class action models. **A)** human and **B)** dog participants. Voxels were ranked according to their feature importances using a random forest classifier, averaged across all iterations of models. The top 5% of voxels (i.e., those used to train models) are presented here, aggregated by species and transformed to group space for visualization purposes (atlases: Mazziotta et al., 2001 for humans and Johnson et al., 2020 for dogs). Labels show dog brain regions with high feature importance scores, based on those identified by Johnson *et al*.^42^; FL is the lateral fissure, FR is the retrosplenial fissure, LPDII is the posterior cingulate gyrus and SSM is the suprasylvian gyrus.

We checked that these species differences weren’t a result of task-correlated motion of the dogs moving more to some types of videos than others (e.g. videos other dogs than, say, cars.) We calculated the Euclidean norm of the six motion parameters and fit a linear mixed-effects model using the R package lme4, with class as a fixed effect and run number as a random effect for each dog. For each of the final models, we found no significant effect of class type on motion for either Daisy (F(2, 2252)=0.83, p=0.44 for object-based and F(4, 1235)=1.87, p=0.11 for action-based) or Bhubo (F(2, 2231)=1.71, p=0.18 for object-based and F(4, 1221)=0.94, p=0.45 for action-based).

## Discussion

Our results demonstrate that naturalistic videos induce representations in dogs’ brains that are stable enough over multiple imaging sessions that they can be decoded with fMRI – similar to results obtained in both humans and monkeys ^20,23^. While previous fMRI studies of the canine visual system have presented stripped-down stimuli, such as a face or object against a neutral background, our results demonstrate that naturalistic videos, with multiple people and objects interacting with each other, induce activation patterns in dog cortex that can be decoded with a reliability approaching that seen in human cortex. This approach opens up new avenues of investigation for how the dog’s visual system is organized.

Although the field of canine fMRI has grown rapidly, to date, these experiments have relied on relatively impoverished stimuli, such as pictures of people, or objects, against neutral backgrounds ^10,11,13^. And while these experiments have begun to identify brain regions analogous to the primate fusiform face area (FFA), involved in face processing, and the lateral occipital cortex (LOC), for object processing, there remains disagreement over the nature of these representations, such as whether dogs have face areas per se, responding to similar salient features as primates, or whether they have separate representations for dogs and humans, or faces and heads, for example ^11,12^. Dogs, of course, are not primates, and we do not know how they interpret these artificial stimuli divorced from their usual multisensory contexts with sounds and smells. Some evidence suggests that dogs do not treat images of objects as representations for the real things ^16^. Although it is not possible to create a true multisensory experience in the scanner, the use of naturalistic videos may mitigate some of the artificialness by providing dynamic stimuli that more closely matches the real-world, at least to a dog. For the same reasons, the use of naturalistic stimuli in human fMRI research has gained popularity, demonstrating, for example, that sequences of events in a movie are represented in the cortex across multiple time scales and that movies are effective at inducing reliable emotion activation ^26^. As such, while naturalistic videos do remain relatively impoverished stimuli, their success in human neuroscience begs the question of whether similar results can be obtained in dogs.

Our results show that a neural net classifier was successful in decoding some types of naturalistic content from dog brains. This success is an impressive feat given the complexity of the stimuli. Importantly, because the classifier was tested on unseen video clips, the decoding model picked up broad categories that were identifiable across clips rather than properties specific to individual scenes. We should note there are multiple metrics for quantifying the performance of a machine-learning classifier (Table 1). Because naturalistic videos, by their nature, will not have equal occurrences of all classes, we took a prudent approach by constructing a null distribution from the random permutation of labels and assessing significance referenced to that. Then, we find that the success of the dog models was statistically significant, achieving 75th-90th percentile scores, but only when the videos were coded based on the actions present, such as playing or talking.

Although the primary goal was to develop a decoder of naturalistic visual stimuli for dogs, comparisons to humans are unavoidable. Here, we note two major differences: for each type of classifier, the human models performed better than dog; and the human models performed well for both object- and action-based models while the dogs performed for action-based only. The superior performance of the human models could be due to several factors. Human brains are roughly ten times larger than dogs’, so there are more voxels from which to choose to build a classifier. To put the models on equal footing, one should use the same number of voxels, but this could be in either an absolute or relative sense. Although the final model was based on the top 5% of informative voxels in each brain (a relative measure), similar results were obtained using a fixed number of voxels. Thus, it seems more likely that performance differences are related to how humans and dogs perceive video stimuli. As noted above, while dogs and humans are both multisensory in their perception, the stimuli may be more impoverished to a dog than a human. Size cues, for example, may be lost, with everything appearing to be a toy version of the real-world. There is some evidence that dogs categorize objects based on size and texture before shape, which is almost opposite to humans ^27^. Additionally, scent, not considered here, is likely an important source of information for object discrimination in dogs, particularly in identification of conspecifics or humans ^28–30^. But even in the absence of size or scent cues, in the unusual environment of the MRI scanner, the fact that the classifier worked at all says that there was still information relevant to the dogs that could be recovered from their brains. With only two dogs and two humans, the species differences could also be due to individual differences. The two dogs, however, represented the best of the MRI-trained dogs and excelled at holding still while watching videos. While a larger sample size would certainly allow more reliable distinctions to be drawn between species, the small number of dogs that are capable of doing awake-fMRI and who will watch videos for periods long enough will always limit generalizability to all dogs. While it is possible that specialized breeds, like sighthounds, might have more finely tuned visual brain responses, we believe that individual temperament and training are more likely to be the major determinants of what is recoverable from a dog’s brain.

These species differences raise the question of what aspect of the videos the dogs were paying attention to. One approach to answering this question relies on simpler video stimuli. Then, by using isolated images of, say, humans, dogs, and cars, both individually and together against neutral backgrounds, we might be able to reverse engineer the salient dimensions to a dog. However, this is both methodologically inefficient and further impoverishes the stimuli from the real-world. The question of attention can be solved by the decoding approach alone, in effect, using the model performance to determine what is being attended to ^31^. Along these lines, our results suggest that while the humans attended to both the actors and the actions, the dogs were more focused on the actions themselves. This might be due to differences in low-level motion features such as the movement frequency when individuals are playing versus eating, or, it might be due to a categorical representation of these activities at a higher level. The distribution of informative voxels throughout the dog’s cortex suggests that these representations are not just low-level features, which would otherwise be confined to visual regions. Further study using a wider variety of video stimuli may illuminate the role of motion in category discrimination by dogs.

In summary, we have demonstrated the feasibility of recovering naturalistic visual information from dog cortex using fMRI in the same way that is done for human cortex. This demonstration shows that even without sound or smells, salient dimensions of complex scenes are encoded by dogs watching videos and that these dimensions can be recovered from their brains. Secondly, based on the small number of dogs that can do this type of task, the information may be more widely distributed in the cortex than typically seen in humans, and the types of actions seem to be more easily recovered than the identity of the actors or objects. These results open up a new way of examining how dogs perceive the environments they share with humans, including video screens, and suggest rich avenues for future exploration of how they, and other non-primate animals, ‘see’ the world.

## Materials and Methods

All methods (both dog and human experiments) were carried out in accordance with relevant guidelines and regulations and reported in accordance with ARRIVE guidelines (https://arriveguidelines.org). The dog study was approved by the Emory University IACUC (PROTO201700572), and all owners gave written consent for their dog’s participation in the study. Human study procedures were approved by the Emory University IRB, and all participants provided written consent before scanning (IRB00069592).

### Participants

Dog participants were two local pet dogs volunteered by their owners for participation in fMRI training and scanning consistent with that previously described ^6^. Bhubo was a 4 year-old male Boxer-mix, and Daisy was an 11 year-old female Boston terrier-mix. Both dogs had previously participated in several fMRI studies (Bhubo: 8 studies, Daisy: 11 studies), some of which involved watching visual stimuli projected onto a screen while in the scanner. They were selected because of their demonstrated ability to stay in the scanner without moving for long periods of time with their owner out of view. Two humans (M, 34yrs and F, 25yrs) also participated in the study. Neither dogs nor humans had previous exposure to the stimuli shown in this study.

### Stimuli

Videos were filmed in Atlanta, Georgia in 2019 using a GoPro HERO7 (1920×1440 pixels, 60fps) mounted on a handheld stabilizing gimbal (Hohem iSteady Pro). Naturalistic videos were filmed from a “dog’s eye view,” holding the gimbal at approximately knee height. Videos were designed to capture everyday scenarios in the life of a dog. These included scenes of walking, feeding, playing, humans interacting (with each other and with dogs), dogs interacting with each other, vehicles in motion, and non-dog animals (Fig. 1A; Movie S1). In some clips, the subjects in the video interacted directly with the camera, for example petting, sniffing or playing with it, while in others, the camera was ignored. Additional footage of deer was obtained from a locally placed camera trap (Bushnell Trophy Cam HD, 1920×1080, 30fps). Videos were edited into 256 unique 7-second ‘scenes’ using Windows Video Editor. Each scene depicted a single event, such as humans hugging, a dog running, or a deer walking. Each scene was assigned a unique number and labelled according to its content (see below).

Scenes were then edited into five larger compilation videos of approximately 6 minutes each. We elected to use compilation videos rather than one long film to allow us to present a wide variety of stimuli in sequence, which would have been difficult to achieve had we attempted to capture them in one long ‘take’. This is consistent with fMRI decoding studies in humans ^20,22^. Additionally, presenting compilations of short clips allowed easier creation of a hold-out set on which the trained algorithm could be tested (see Analyses below), as we were able to hold out individual clips instead of one long movie. Four compilation videos had 51 unique scenes and one had 52. There were no breaks or blank screens between scenes. Scenes were selected semi-randomly to ensure that each video contained exemplars from all the major label categories – dogs, humans, vehicles, non-human animals and interactions. During the compiling process all scenes were downsampled to 1920×1080 pixels at 30 frames per second to match the resolution of the MRI-projector.

### Experimental Design

Participants were scanned in a Siemens 3T Trio MRI scanner while watching the compilation videos projected onto a screen mounted at the rear of the MRI bore. Videos were played without sound. For dogs, stable positioning of the head was achieved by prior training to place their head in a custom-made chin rest, molded to the lower jaw from mid-snout to behind the mandible. The chin rest was affixed to a wood shelf that spanned the coil but allowed enough space for the paws underneath. The result was each dog assuming a ‘sphinx’ position (Fig. 1B). No restraints were used. For further information on training protocol, see previous awake fMRI dog studies ^6^. Subjects participated in 5 runs per session, where each run consisted of one compilation video watched start to finish, presented in a random order. For dogs, short breaks were taken between each run, during which food rewards were delivered to the dog. Each subject participated in 3 sessions over two weeks and so watched each of the 5 unique compilation videos 3 times, yielding an aggregate fMRI time of 90 minutes per individual.

### Imaging

Dog participants were scanned following a protocol consistent with that employed in previous awake fMRI dog studies ^6,32^. The functional scans were obtained using a single-shot echo-planar imaging sequence to acquire volumes of 22 sequential 2.5 mm slices with a 20% gap (TE = 28 ms, TR = 1430 ms, flip angle = 70°, 64 × 64 matrix, 2.5 mm in-plane voxel size, FOV = 160 mm). For dogs, slices were oriented dorsally to the brain with the phase-encoding direction right-to-left, as dogs sit in the MRI in a ‘sphinx’ position, with the neck in line with the brain. Phase encoding right-to-left avoids wrap-around artifacts from the neck into the front of the head. In addition, the major susceptibility artifact in scanning dogs comes from the frontal sinus, resulting in distortion of the frontal lobe. We find right-to-left distortion preferable to stretching or compressing the frontal lobe. For humans, axial slices were obtained with phase-encoding in the anterior-posterior direction. To allow for comparison with the dog scans (same TR/TE), multiband slice acquisition was used (CMRR, University of Minnesota) for the humans with a multiband acceleration factor of 2 (GRAPPA=2, TE=28 ms, TR=1430 ms, flip angle = 55°, 88 × 88 matrix, 2.5 mm in-plane voxels, 44 2.5 mm slices with a 20% gap). For the dogs, a T2-weighted structural image of the whole brain was also acquired for each participant using a turbo spin-echo sequence with 1.5mm isotropic voxels. For the humans, a T1-weighted MPRAGE sequence with 1 mm isotropic voxels was used. Over the course of three sessions, approximately 4000 functional volumes were obtained for each participant.

### Stimulus Labels

In order to train a model to classify the content presented in the videos, the scenes first had to be labelled. To do this, we divided the 7s scenes that made up each compilation video into 1.4 s clips. We chose to label short clips rather than individual frames as there are elements of video that cannot be captured by still frames, some of which may be particularly salient to dogs and therefore useful in decoding, such as movement. We chose a clip length of 1.4 s because this was long enough to capture these dynamic elements and closely matched the TR of 1.43 s, allowing us to perform the classification on a volume-by-volume basis. These 1.4 s clips (n=1280) were then randomly distributed to lab members who manually labelled each clip using a pre-programmed check-box style submission form. There were 94 labels, chosen to encompass as many key features of the videos as possible, including subjects (e.g., dog, human, cat), number of subjects (1, 2, 3+), objects (e.g., car, bike, toy), actions (e.g., eating, sniffing, talking), interactions (e.g., human-human, human-dog) and setting (indoors, outdoors), among others. This produced a 94-dimensional label vector for each clip (Dataset S1). As a consistency check, a random subset was selected for relabeling by a second lab member, and labels were found to be highly consistent across individuals (>95%). For those labels that were not consistent, the two lab members rewatched the clip in question and together came to a consensus on label.

For each run, timestamped log files were used to determine the onset of the video stimulus relative to the first scan volume. To account for the delay between stimulus presentation and the BOLD response, labels were then convolved with a double gamma hemodynamic response function (HRF) and interpolated to the TR of the functional images (1430 ms) using the Python functions numpy.convolve() and interp(). The end result was a matrix of convolved labels by the total number of scan volumes for each participant (94 labels x 3932, 3920, 3939 and 3925 volumes for Daisy, Bhubo, Human 1 and Human 2 respectively). These labels were later grouped where necessary to create macrolabels for further analysis. For example, all instances of walking (dog walking, human walking, donkey walking) were combined to create a ‘walking’ label. To further remove redundancy in the label set, we calculated the Variance Inflation Factor (VIF) for each label, excluding the macrolabels, which were obviously highly correlated. VIF is a measure of multicollinearity in predictor variables, calculated by regressing each predictor against every other. Higher VIFs indicate more highly correlated predictors. We employed a VIF threshold of 2, reducing the 94 labels to 52 unique, largely uncorrelated labels (Dataset S1).

### fMRI Preprocessing

Preprocessing involved motion correction, censoring and normalization using the AFNI suite (NIH) and its associated functions ^33,34^. Two-pass, six-parameter rigid-body motion correction was used to align volumes to a target volume that was representative of the participant’s average head position across runs. Censoring was performed to remove volumes with more than 1 mm displacement between scans, as well as those with outlier voxel signal intensities greater than 0.1%. For both dogs, more than 80% of volumes were retained after censoring, and for humans, more than 90% were retained. To improve the signal-to-noise ratio of individual voxels, mild spatial smoothing was performed using 3dmerge and a 4 mm Gaussian kernel at full-width half-maximum.

To control for the effect of low level visual features such as motion or speed that may differ according to stimulus, we calculated the optical flow between consecutive frames of video clips ^22,35^. Optical flow was calculated using the Farneback algorithm in OpenCV, after down sampling to 10 frames per second ^36^. To estimate the motion energy in each frame, we calculated the sum of squares of the optic flow of each pixel and took the square root of the result, effectively calculating the Euclidean average optic flow from one frame to the next ^35,37^. This generated time courses of motion energy for each compilation video. These were resampled to match the temporal resolution of the fMRI data, convolved with a double gamma hemodynamic response function (HRF) as above, and concatenated to align with stimulus presentation for each subject. This time course, along with the motion parameters generated from the motion correction described above, were used as the only regressors to a general linear model (GLM) estimated for each voxel with AFNI’s 3ddeconvolve. The residuals of this model were then used as inputs to the machine-learning algorithm described below.

### Analyses

We aimed to decode those regions of the brain that contribute significantly to classification of visual stimuli, training a model for each individual participant that could then be used to classify video content based on participants’ brain data. We used the Ivis machine learning algorithm, a nonlinear method based on Siamese neural networks (SNNs) that has shown success on high dimensional biological data ^38^. SNNs contain two identical sub-networks that are used to learn the similarity of inputs in either supervised or unsupervised modes. Although neural networks have grown in popularity for brain decoding because of their generally greater power over linear methods like support vector machines (SVMs), we used an SNN here because of its robustness to class imbalance and need for fewer exemplars. Compared to support vector machines (SVM) and random forest (RF) classifiers trained on the same data, we found Ivis to be more successful in classifying brain data across multiple label combinations, as determined by various metrics including mean F1 score, precision, recall and test accuracy (see below).

For each participant, the whole-brain residuals were converted to a format appropriate for input into the Ivis neural network. The five runs in each of their three sessions were concatenated and masked, retaining only brain voxels. The spatial dimension was then flattened, resulting in a two-dimensional matrix of voxels by time. The convolved labels of the videos shown in each run were also concatenated, thus corresponding to the fMRI runs. Both the fMRI data and corresponding labels were censored according to the volumes flagged in preprocessing. Target labels to be decoded – hereafter referred to as ‘classes’ - were selected and only those volumes containing these classes were retained. For simplicity, classes were treated as mutually exclusive, and volumes belonging to multiple classes were not included for decoding, leaving only pure exemplars. The data were split into training and test sets. We used a 5-fold split, randomly selecting 20% of the scenes to act as the test set. This meant that if a given scene was selected for the test set, all the clips and functional volumes obtained during this scene were held out from the training set. Had the split been performed independent of scene, volumes from the same scene would appear in both the training set and the test set, and the classifier would only have to match them to that particular scene to be successful in classifying them. However, to correctly classify held-out volumes from new scenes, the classifier would have to match them to a more general, scene-independent class. This was a more robust test of the generalizability of the classifier’s success compared to holding out individual clips. The training set was then balanced by undersampling the number of volumes in each class to match that of the smallest class using the scikit-learn package imbalanced-learn.

For each participant, we trained and tested the Ivis algorithm on 100 iterations, each time using a unique test-train split (Ivis parameters: k=5, model=’maaten’, n_epochs_without_progress=30, supervision_weight=1). These parameter values were largely selected on the basis of dataset size and complexity as recommended by the algorithm’s authors in its documentation ^39^. ‘Number of epochs without progress’ and ‘supervision weight’ (0 for unsupervised, 1 for supervised) underwent additional parameter tuning to optimize the model. To reduce the number of features used to train the classifier from the whole brain to only the most informative voxels, we used a Random Forest Classifier (RFC) using scikit-learn to rank each voxel according to its feature importance. Although the RFC did not perform above chance by itself, it did serve the useful purpose of screening out noninformative voxels, which would have contributed only noise to the Ivis algorithm. This is similar to using F-tests for feature selection before passing to the classifier ^40^. Only the top 5% of voxels from the training set were used in training and testing. 5% was selected as the preferred number of voxels as a conservative threshold in an effort reduce the number of noninformative voxels prior to training the neural net. Qualitatively similar results were also obtained for both humans and dogs when using a larger proportion of voxels. Though human brains are larger than dog brains, human models were also successful when trained on an absolute number of voxels equal to those included in dog models, far smaller than 5% of voxels (∼250 voxels; all mean LRAP scores > 99th percentile). For consistency, we therefore present results using the top 5% of voxels for both species. The average 5% most informative voxels across all 100 runs were then normalized, transformed to each participant’s structural space and then to group atlas space (atlases: Mazziotta et al., 2001 for humans and Johnson et al., 2020 for dogs) and summed across participants for each species. Feature importances were overlayed on atlases and colored according importance score using ITK-SNAP ^43^.

The output layer of the Ivis network contained a number of elements equal to the number of training classes. The most common metrics to assess model performance in machine learning analyses include precision, accuracy, recall and F1 score. Accuracy is the overall percentage of model predictions that were correct, given the true data. Precision is the percentage of the model’s positive predictions that are actually positive (i.e. the true positive rate), while recall is the percentage of true positives in the original data that the model was able to successfully predict. F1 score is the weighted average of precision and recall and acts as an alternate measure of accuracy that is more robust to class imbalance. However, the Ivis differs from other commonly used machine learning algorithms in that its output is not binary. Given a particular input of brain voxels, each output element represented the probabilities corresponding to each of the classes. Computing accuracy, precision, recall and F1 for these outputs required binarizing them in a ‘winner takes all’ fashion, where the class with the highest probability was considered the one predicted for that volume. This approach would eliminate important information about the ranking of these probabilities that was relevant to assessing the quality of the model. Thus, while we still computed these traditional metrics, we used the Label Ranking Average Precision (LRAP) score as the primary metric to compute the accuracy of the model on the test set. This metric essentially measures to what extent the classifier assigned higher probabilities to true labels ^25^.

Test sets, unlike the training sets, were not balanced across classes. Comprising only 20% of the data, undersampling to the smallest class size would have resulted in very small sample sizes for each class, such that any statistics calculated would have been unreliable. To avoid the possibility of inflated accuracy from this imbalance, we computed the null distribution of the LRAP by randomly permuting the order of the classes 1000 times for each model iteration. This null distribution acted as a reference for how well the model was likely to perform by chance. The true LRAP was then converted to a percentile ranking in this null distribution. A very high percentile ranking, for example 95%, would indicate that a score that high arose only 5% of the time in 1000 random permutations. Such a model could therefore be deemed to be performing well above chance. To determine if these percentile rankings were significantly greater than that expected by chance – that is, the 50th percentile – statistically, we calculated the median LRAP percentile ranking across all 100 iterations for each model and performed a one-sample Wilcoxon signed rank test.

We carried out a series of preliminary exploratory analyses to determine which classes to include in our models. This included computing dissimilarity matrices for the entire 52 potential classes of interest using the Python package scipy’s hierarchical clustering algorithm, which clustered classes based on the similarity of an individual’s brain response to each as defined by pairwise correlation. The resulting clusters suggested that classes denoting similar actions, for example human eating and dog eating, may have evoked a more similar brain responses in dogs than those with the same subject, for example human eating and human walking. This prompted us to explore action versus subject-based models, as well as models attempting to decode both, with a variety of class combinations from the classes with sufficient data to include. The models presented here are the most successful and scientifically relevant of those trials.

## Data Availability

All study data, including videos, video labels, fMRI data and Python code used for analysis, available at datadryad.org (url tk).

## Acknowledgments

We thank Kate Revill, Raveena Chhibber and Jon King for their helpful insights in the development of this analysis, Mark Spivak for his assistance recruiting and training dogs for MRI, and Phyllis Guo for her help in video creation and labelling. We also thank our dedicated dog owners, Rebecca Beasley (Daisy) and Ashwin Sakhardande (Bhubo). Human studies supported by a grant from the National Eye Institute (Grant R01 EY029724 to D.D.D.)

## Author Contributions

E.M.P. and G.S.B. designed research; E.M.P., K.D.G., D.D.D. and G.S.B. performed research; E.M.P. analyzed data; and E.M.P. and G.S.B. wrote the paper.

